# Experimental transmission of African swine fever virus by *Ornithodoros lahorensis* (Acari: Argasidae)

**DOI:** 10.1101/2025.03.24.644965

**Authors:** Jin Luo, Huaijie Jia, Shuaiyang Zhao, Qiaoyun Ren, Muhammad Kashif Obaid, Jifei Yang, Guiquan Guan, Hong Yin, Guangyuan Liu, Qingli Niu

**Author notes:** E-mail addresses: JL, HJJ, SYZ, QYR, MKO, JFY, GQG, HY, GYL, QLN.

## Abstract

African swine fever (ASF) is an acute, highly contagious infectious disease that affects both domestic and wild pigs, caused by the African swine fever virus (ASFV). ASFV is the only known DNA virus transmitted by arthropod vectors, with acute infection in pigs leading to morbidity and mortality rates as high as 100%.The virus can persist in a transmission cycle among wild boars, soft ticks, and domestic pigs. To date, nine *Ornithodoros* spp. have been confirmed to be capable of transmitting ASFV worldwide. However, the potential soft tick species capable of transmitting ASFV in China remain unclear. In this study, we compared the ability of *Argasidae* spp. *Ornithodoros lahorensis* and *Argas persicus*, as well as *Ixodidae* spp., *Haemaphysalis longicornis*, *Rhipicephalus sanguineus* and *Dermacentor silvarum,* to transmit ASFV via animal transmission experiments. The results revealed that *O. lahorensis* soft ticks, but not *A. persicus* or Ixodes, could act as competent vectors through transstadial and transovarial transmission of ASFV. The virus titers horizontally transmitted ASFV to ticks were 10^5.83^, 10^6.59^ and 10^4.31^, HAD50/ml respectively. These viruses were detected in the nymphs developed from larvae, adults developed from nymphs, and larvae hatched from eggs by adults. Thus, *O. lahorensis* ticks are likely an important natural vectors of ASFV, although both mammalian and tick hosts are likely required for the maintenance of ASFV in the sylvatic cycle.

## Introduction

Ticks are major ectoparasites causing significant economic losses and serving as vectors for livestock diseases ^1^. They are classified into two families: Ixodidae (hard ticks) with over 879 species, and Argasidae (soft ticks) with approximately 200 species worldwide ^2^. Tick competence refers to their ability to acquire, maintain, multiply, and transmit pathogens to new hosts. This process involves four stages: ingestion of an infected blood meal, pathogen infection in the midgut, persistence and multiplication despite the tick’s immune response, and dissemination to organs like the salivary glands for transmission ^3,4^. The distribution and prevalence of ticks are influenced by host availability, tick populations, and environmental conditions ^5^.

African swine fever virus (ASFV) is a highly contagious and lethal virus that causes African swine fever (ASF), an acute hemorrhagic fever in both domestic and wild pigs. ASFV is the only known DNA virus transmitted by arthropod vectors, particularly soft ticks from the Ornithodoros genus (Argasidae family) ^6–8^. The virus is characterized by a complex genome, containing a large, double-stranded DNA sequence that encodes numerous structural and non-structural proteins involved in immune evasion and host cell modulation ^9^. ASFV causes high morbidity and mortality in infected pigs, with mortality rates reaching 100% in some outbreaks, leading to severe economic losses in the global swine industry ^10^.

ASFV transmission occurs through direct contact with infected pigs or via tick vectors. Soft ticks, such as *O. moubata*, play a key role in the transmission cycle, allowing the virus to persist in wildlife reservoirs like wild boar and certain tick populations ^11^. ASFV has no known vaccine or effective treatment, making its control challenging. Efforts to understand ASFV pathogenesis, transmission dynamics, and vector competence are critical for developing strategies to prevent and control ASF outbreaks. The recent resurgence of ASF outbreaks has highlighted the urgent need for understanding the mechanisms by which this virus is transmitted ^12^. Vector-borne diseases often involve intricate interactions between pathogens and their vectors, complicating the dynamics of transmission. The ability of a vector to transmit a pathogen is influenced by several biological and ecological factors, including the host’s immunological response, environmental conditions, and biological characteristics, such as feeding behavior and lifespan ^13,14^. ASFV has been detected in 9 species of soft ticks: *O. turicata* ^15^, *O. phacochoerus*, *O. porcinus* ^16^, *O. erraticus*, *O. moubata* ^17^, *O. waterbergensis*, *O. compactus* ^18^, *O. papillipes* ^19^ and *O. verrucosus* ^20^.

*O. lahorensis* is a species of soft tick; compared with the ixodid group, argasid ticks are nidicolous or endophilic and prefer to inhabit sheltered areas such as caves, nests, and burrows of animal hosts, as well as human homes. Soft ticks usually spend 15–30 min per feeding event ^21^ and feed in multiple life stages. Soft ticks can transmit pathogens from their midgut into the host via saliva secretions ^22^. *O. lahorensis* is an important pathogenic vector, and with the movement of its hosts, this tick has become distributed worldwide and is considered a reservoir for several diseases ^23^. However, because of the short feeding duration and nidicolous lifestyle, fewer studies have been conducted on argasid ticks and argasid tick-borne diseases than on ixodid ticks. Research has shown that *O. lahorensis* can also transmit *Babesia* spp., *Theileria* spp., *Anaplasma ovis* ^24,25^, and *Borrelia* spp. ^26^. However, it is unknown whether *O. lahorensis* can transmit ASFV.

To date, three reported avian tick species in China, *O. lahorensis*, *O. tartaki*, and *O. papillipes*, have been reported to bite humans ^27^. No reports have been published on the transmission of ASFV by these important vector species. *O. lahorensis* is the most common of these species and, in China, the most harmful. In this study, we investigated the ability of *O. lahorensis* to transmit ASFV within the framework of animal experiments. The findings can help reveal the transmission patterns of this disease within ecosystems and provide a theoretical basis for effective epidemic control.

## Materials and Methods

### Ethics approval

All ASFV studies were conducted at the biosecurity level 3 (BSL-3) laboratory of the Lanzhou Veterinary Research Institute (LVRI), Chinese Academy of Agricultural Sciences, accredited by the China National Accreditation Service for Conformity Assessment (CNAS) and approved by the Ministry of Agriculture and Rural Affairs. To mitigate potential risks, strict adherence to protocols was required, with all activities monitored by qualified LVRI personnel and local and central government authorities conducting unannounced inspections. All animals were handled according to the Animal Ethics Procedures and Guidelines of the People’s Republic of China. This study was approved by the Animal Administration and Ethics Committee of Lanzhou Veterinary Research Institute, Chinese Academy of Agricultural Sciences (approval No. LVRIAEC 2024-006).

### Ticks and virus

In this study, *O. lahorensis* and *Argas persicus* soft ticks were collected from Gaotai County in China, *Haemaphysalis longicornis* and *Dermacentor silvarum* from Qingyang city in Gansu Province, and *Rhipicephalus sanguineus* from Xinjiang Ili Kazakh Autonomous Prefecture. The ticks were collected in transparent glass tubes with open ends, which were then sealed with breathable white cloth to prevent their escape. The collected ticks were taken back to the laboratory for artificial cultivation. ASFV CN/SC/2019 was isolated, identified, and maintained in LVRI-BSL-3 of the African Swine Fever Regional Laboratory of China (Lanzhou).

### RT‒qPCR

The B646L gene was amplified from field-collected tick samples using the following ASFV-specific primers: F: 5′-CT TGC GAT CTG GAT TAA GCT GCG C-3′; R: 5′-CAC AAG ATC AGCT AGT GAT AG-3′ ^28^. For PCR, 5 μL of 10× buffer, 5 μL of dNTPs (200 μmol/L), 1 μL of each of the F/R primers (10–100 μmol/L), 1 μL of Taq DNA polymerase (2.5 U), and 2 μL of gDNA template (1–2 ng/μL) were added to the PCR tube, and double distilled water was added to bring the total volume to 50 μL. After mixing amplification was performed with the following program: 97°C for 5 min; 40 cycles of 95°C for 40 s, 56°C for 30 s, and 72°C for 1 min; and 73°C for 5 min. The PCR products were subjected to electrophoresis.

### Animal transmission experiments

All the collected ticks were identified at the species level using taxonomic keys and morphological characteristics ^29^. To confirm whether *O. lahorensis* is a vector for ASFV transmission. The ASFV strains were inoculated into clean, healthy pigs. After infection, laboratory-reared clean hungry *O. lahorensis* larvae, nymphs, and adults were released onto the infected pig and secured with a cloth bag to ensure that the ticks could not escape from the pig. Soft ticks quickly feed on blood, and switch hosts or continue to feed on the original host until they become engorged, and develop to the next stage. In this process, the nymph stage of *O. lahorensis* involves four developmental stages: stages I to II. During the blood-feeding process, the nymphs do not become engorged on the host, but they do molt. The whole process involves three in which the nymph does not leave the host. After the fourth stage of full feeding, the nymphs fall off and molt into hungry adults. The entire development process lasts 27 ± 5 days. In the experiment, we collected fully fed larvae, stage IV fully fed nymphs, and engorged adults, which were then maintained at room temperature (23°C) and a relative humidity of 87± 5%. The strength of ASFV infection was determined via qPCR.

On the other hand, to verify that ASFV can be transmitted through the eggs of *O. lahorensis*, ticks that had fed on the blood of ASFV-infected pigs in the laboratory were allowed to lay eggs, which were then hatched. ASFV infection in hatched larvae was detected by qPCR, and some of the larvae were released onto the surface of another clean, ASFV-free pig. The clinical symptoms of the pigs were observed daily, and after approximately 5‒7 days, the pigs presented typical symptoms of ASFV infection, such as high fever, anorexia, skin erythema, and, in the later stages of the disease, bloody stools and bleeding from the mouth and nose. The experimental pigs died approximately 15 days later. During the above experimental process, blood samples from the pigs during the infection period and tissue samples from the dead pigs were collected and tested by qPCR.

### Data analysis

Molecular detection methods were used to determine the presence of ASFV within the field-collected ticks and infected ticks used for transmission experiments. In this process, total gDNA was extracted from different tick samples according to the manufacturer’s instructions for standard DNA reagents (Invitrogen, 74016) and used as a template for amplification analysis of target genes. qPCR experiments were performed according to the manufacturer’s instructions for SYBR Green (Bio-Rad, 1725124) to quantify target gene expression. The *β*-actin gene was used as an internal reference control, and the sequences of the primers used were as follows: F: 5’-TGT GAC GAC GAG GTT GCC G-3’; R: 5’-GAA GCA CTT GAG GTG GAC AAT G-3’ ^30^. The relative expression of each designated gene was calculated by the 2^−ΔΔCt^ method.

### Transmission electron microscopy observation

The ticks were subjected to cold treatment, and once immobilized, they were subjected to dissection using sterile instruments to isolate the salivary glands. This task was performed under a stereomicroscope to ensure precision and minimize contamination. The dissected tissues were then fixed in a suitable chemical fixative, such as glutaraldehyde or paraformaldehyde, which preserved the cellular structure and integrity of the viruses present. The samples were subsequently subjected to dehydration through an ethanol gradients to prepare them for embedding in a resin, a critical step for achieving the requisite mechanical stability for PEM analysis.

Following dehydration, the tissues were embedded in epoxy resin. The resin offered the rigidity necessary for thin sectioning, which is indispensable for electron microscopy. Once embedding was completed, the samples were polymerized, and ultrathin 70–90-nanometer-thick sections were cut with an ultramicrotome. The choice of section thickness is crucial, as the thickness affects the visibility of the structures within the tissues and ensures that individual viral particles are distinguishable.

## Results

### Morphological characteristics of *O. lahorensis*

*O. lahorensis* ticks are relatively large, with adult body lengths reaching up to 10 millimeters. They exhibit a distinctly flattened shape, with a rounded dorsal surface that appears smooth and is brown or black in color. The frontal region is relatively broad, with a straight anterior edge and a slightly curved posterior edge. The mouthparts of this tick species are quite prominent and appear more pronounced than those of other tick species. The dorsal side of the *O. lahorensis* body features a well-known reticulated pattern composed of geometric spots, and the arrangement of these spots is characteristic of the variety. The color of these spots’ ranges from light brown to dark brown, facilitating camouflage (Figure 1A). Additionally, the ventral side displays significant copulatory structures; the male tick’s copulatory organ is typically “Y”-shaped, whereas the female exhibits a more complex structure (Figure 1B).

**Figure 1.**
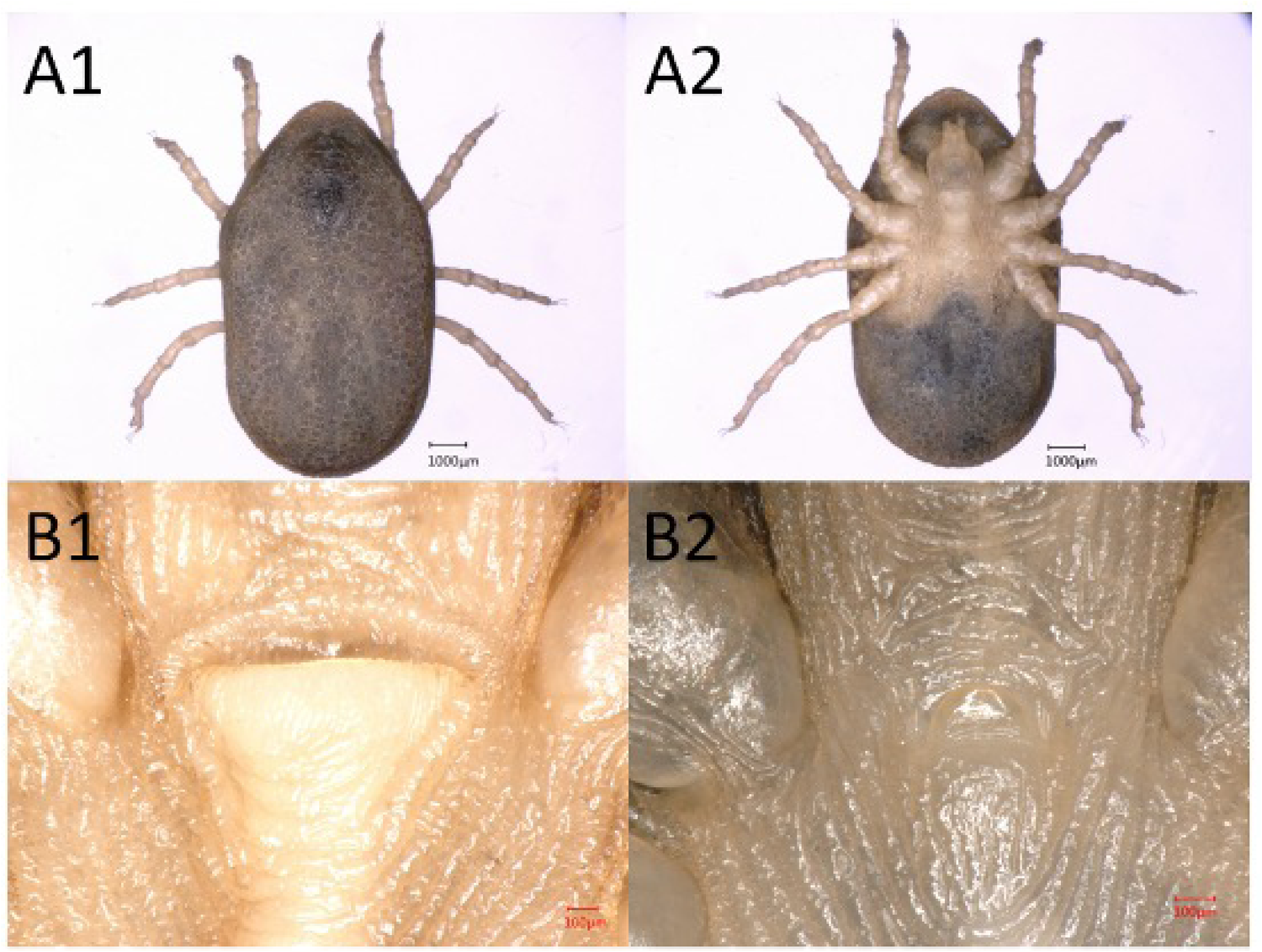
Morphological characteristics of *O. lahorensis*. **A1-A2**: Dorsal and ventral views of the morphology of *O. lahorensis*, respectively; **B1-B2**: genital pores of female and male *O. lahorensis,* respectively

### Detection of ASFV in wild ticks

#### ASFV has not been transmitted to ticks in China

In this study, ASFV nucleic acids were detected in *O. lahorensis*, *Argas persicus* soft ticks, *Haemaphysalis longicornis*, *Dermacentor silvarum* and *Rhipicephalus sanguineus* collected from different regions of Northwest China. However, all the results were negative (data not shown).

#### Transmission of ASFV by different tick species to susceptible domestic pigs

A transmission experiment was performed to confirm whether these ticks can transmit ASFV. However, except *O. lahorensis,* all the ticks that fed on ASFV-infected pigs tested negative by PCR when they developed to the next stage in an unengorged state (Table 1). Moreover, when the infected ASFV larvae developed into unengorged nymphs, the nymphs were allowed to feed on three clean pigs, which resulted the death of two pigs with an ASFV titer of 10^5.83^ HAD50/ml in the blood of pig. Similarly, when infected ASFV nymphs developed into unengorged adults and fed on clean pigs, all three pigs died. The ASFV titers in the pig’s blood transmitted by adults the highest, reaching 10^6.59^ HAD50/ml. After the infected ASFV *O. lahorensis* adults laid eggs the hatched larvae were allowed to feed on three clean pigs, which resulted in the death of two pigs, and the virus titer in the blood of the dead was 10^4.31^ HAD50/ml. The same experiment was conducted with representative strains of different genera, including *Argas persicus*, *Haemaphysalis longicornis*, *Dermacentor silvarum*, and *Rhipicephalus sanguineus*.

**Table 1.**
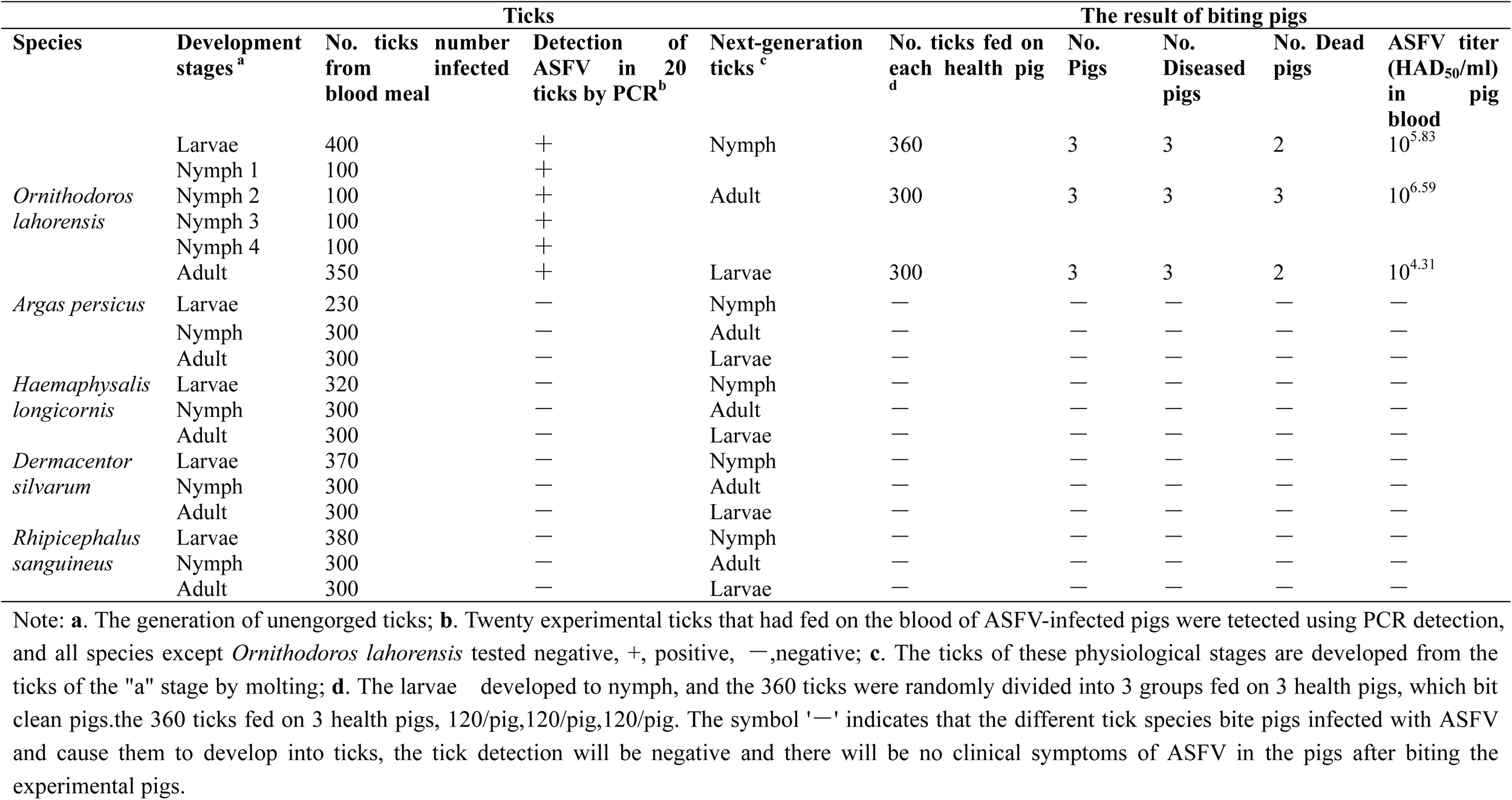
Transmission of African swine fever virus by different tick species to susceptible domestic pigs.

#### ASFV successfully transmited to next generation of *O. lahorensis*

The results of the ASFV transmission experiment showed that after the unengorged nymphs, unengorged larvae, and unengorged adults fed on pigs infected ASFV, they were allowed to develop to the next physiological stage, i.e., hungry nymphs, hungry adults, and egg laying. These developed ticks were taken ASFV detection. It was shown that these developed ticks all carried the ASFV. To confirm that these ticks could transmit the virus to other clean pigs, this released these ASFV-positive unengorged ticks onto the surface of another clean pig to feed. During this process, pig blood was collected daily, and PCR detection was performed on swabs, rectal swabs, and oral swabs of the bitten pigs. About 7 days later, all the above swabs were able to detect positive. These results confirmed that *O. lahorensis* could can transmit ASFV period (horizontally) (Figure 2). In addition, after the infected adult ticks eggs, the eggs were incubated for nearly a month, and the unengorged larvae were detected for their carrying of ASFV, and at the same time, they were onto the surface of a clean pig to feed, and blood was collected daily and stored at -80 ℃ for later use. This experiment confirmed that *O. lahorensis* could transmit ASFV through eggs (Figure 2).

**Figure 2.**
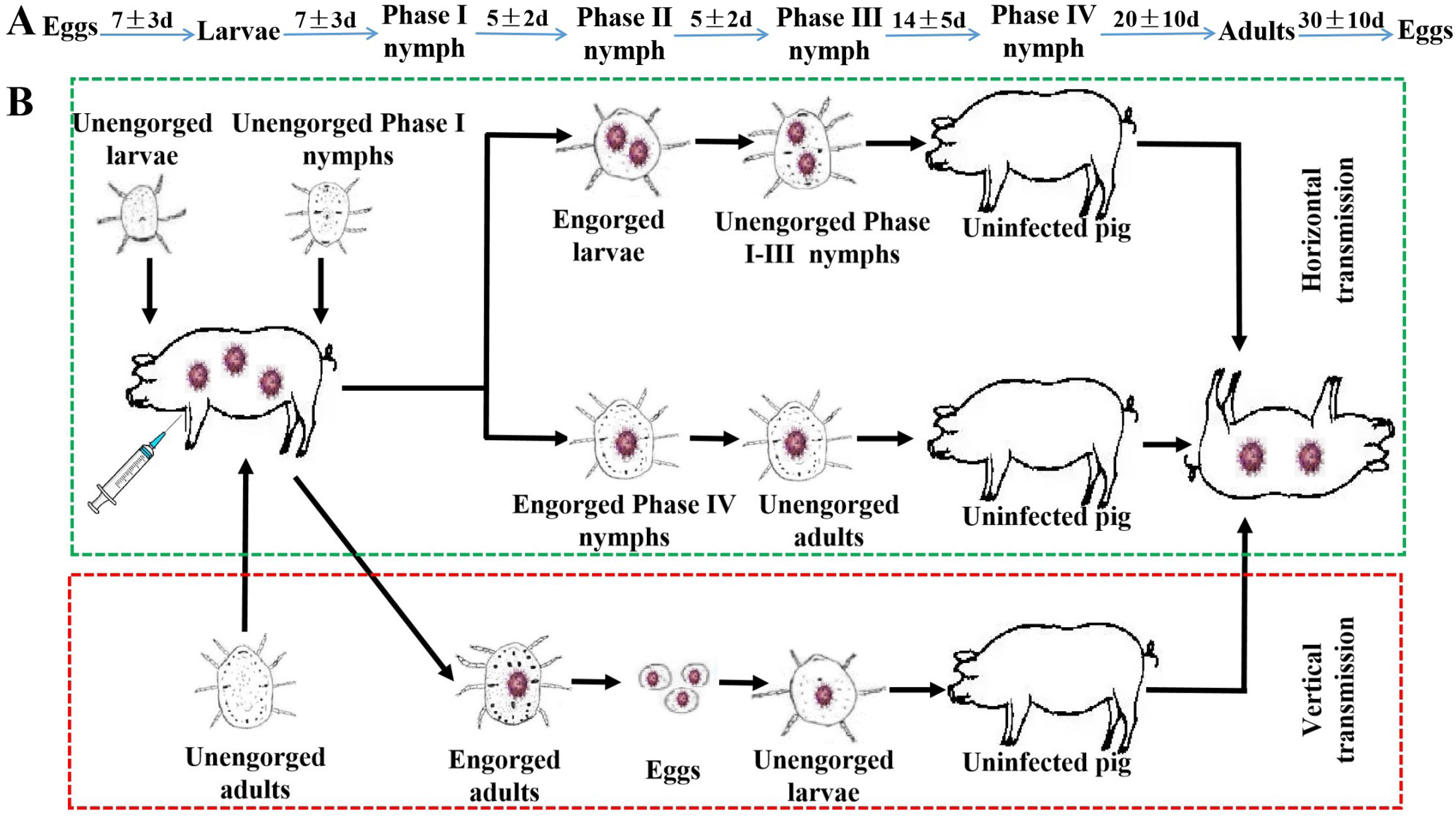
Schematic diagram of *O. lahorensis* as a biological vector for ASFV. **A.** Characteristics of the maintenance system of *O. lahorensis* in laboratory conditions; **B.** The green box indicates that ASFV can be transmitted horizontally in *O. lahorensis*; the red box indicates that ASFV can be transmitted vertically in *O. lahorensis*.

#### Effect of ASFV infection on tick development

Whether the ASFV infection can cause changes in the normal physiological development of the Lahore soft tick is still unknown. Our study showed that the ticks (n=80) in the infected Group1 laid eggs, with one tick laying only a small number of eggs and then dying, the egg laying was terminated. Approximately 21,000 eggs were laid by 70 ticks. In the non-infected Group2, the 74 of the 80 ticks laid eggs, with 70 ticks laying about 23,000 eggs. The hatching rate of the in the experimental group was 80.96%, and that of the control group was 76.09% (Table 2). In addition, also counted the engorged weight and mortality rate of the ticks. The results showed that the engorged weight of the larvae, nymphs, and adults infected ASFV were 8.83 ± 1 (n=50), 81.72±6 (n=10), and 13.48±10 (n=10), respectively. These values were not significantly different from those of the control group (p-value of 0176, 0.312, and 0.059, respectively). There was also no significant difference in the mortality rate. The mortality of the larvae was higher, with 36.75% in the experimental group and 32.75% in the control group; the mortality of the nymphs was 23.50% in the experimental group and 10.50% in the control group; the mortality rate of adults was 14.00% in the experimental group and 18.00% in the control group (Table 3). These results indicate the infection of ASFV does not affect the normal physiological development of *O. lahorensis*.

**Table 2.** The oviposition range (%) of *O. lahorensis* females during ASFV infection.

| Group | No. ticks on each pig | Oviposition rate (%) | Numbers of Eggs | Hatched rate (%) |
| --- | --- | --- | --- | --- |
| 1 | 80 | 71<sup>a</sup>/80<sup>b</sup> (88.75%) | 21000 (70<sup>c</sup>) | 17000<sup>d</sup>/21000<sup>e</sup> (80.95%) |
| 2 | 80 | 74/80 (92.50%) | 23000 (70) | 17500 /23000 (76.09%) |

Note: Group 1, pigs infected with ASFV; Group 2, pigs did not infect with ASFV. a/b: a, number of females that oviposited; b, number of females that fed. c, Number of ticks that laid eggs. d/e: d, number of hatched eggs; e, total number of eggs produced by all ticks.

**Table 3.** Weights (mg) of *O. lahorensis* larvae, nymphs, and adults immediately after feeding on pigs infected with ASFV.

| Group | Stages | No. ticks | Engorged weight, mean (mg)<br>$\bar{X} \pm \text{SD}$ | p-value | Mortality rate (%) |
| --- | --- | --- | --- | --- | --- |
| 1 | Larvae | 400 | 8.83±1 (50<sup>a</sup>) | 0.176 | 147/400 (36.75%) |
|  | Nymphs | 200 | 81.72±6 (10<sup>b</sup>) | 0.312 | 47/200 (23.50%) |
|  | Adults | 100 | 103.48±10 (10<sup>c</sup>) | 0.059 | 14/100 (14.00%) |
| 2 | Larvae | 400 | 10.17±3 (50) | — | 131/400 (32.75%) |
|  | Nymphs | 200 | 76.06±6 (10) | — | 21/200 (10.50%) |
|  | Adults | 100 | 120.53±8 (10) | — | 18/100 (18.00%) |

Note: Group 1, ticks infected with ASFV; Group 2, ticks did not infect with ASFV. a, average weight of 50 engorged larvae; b, average weight of 10 engorged nymphs; c, average weight of 10 engorged adults.

#### ASFV could replication in tick tissues

In this study, we used transmission transmission electron microscopy (TEM) and PCR methods to detect ASFV in different anatomical tissues of *O. lahorensis*. The results showed that ASFV particles were observed in the salivary gland traces at 15 days later, and were accompanied by significant rough endoplasm reticulum proliferation (Figure 3A). In Figure 3B, it can be observed that the mitochondria in the salivary gland cells gradually expand which provides energy for the maturation of the viral factory ASFV. To further confirm that *O. lahorensis* is a biological vector of ASFV, The gDNA from different tissues (salivary glands, midgut, Malpighian tubules, basi, epidermis, and fibrous tissue) of the ticks and performed PCR amplification of B646L (uninfected ticks were used as controls). The results showed that there a certain degree of infection in various tissues of *O. lahorensis* infected with ASFV (Figure 3C). The expression of the virus in these tissues was quantified, indicating that the content of the virus in the tick, with the development of the tick or the maturity of various organs, the content of the virus was (the results were not shown).

**Figure 3.**
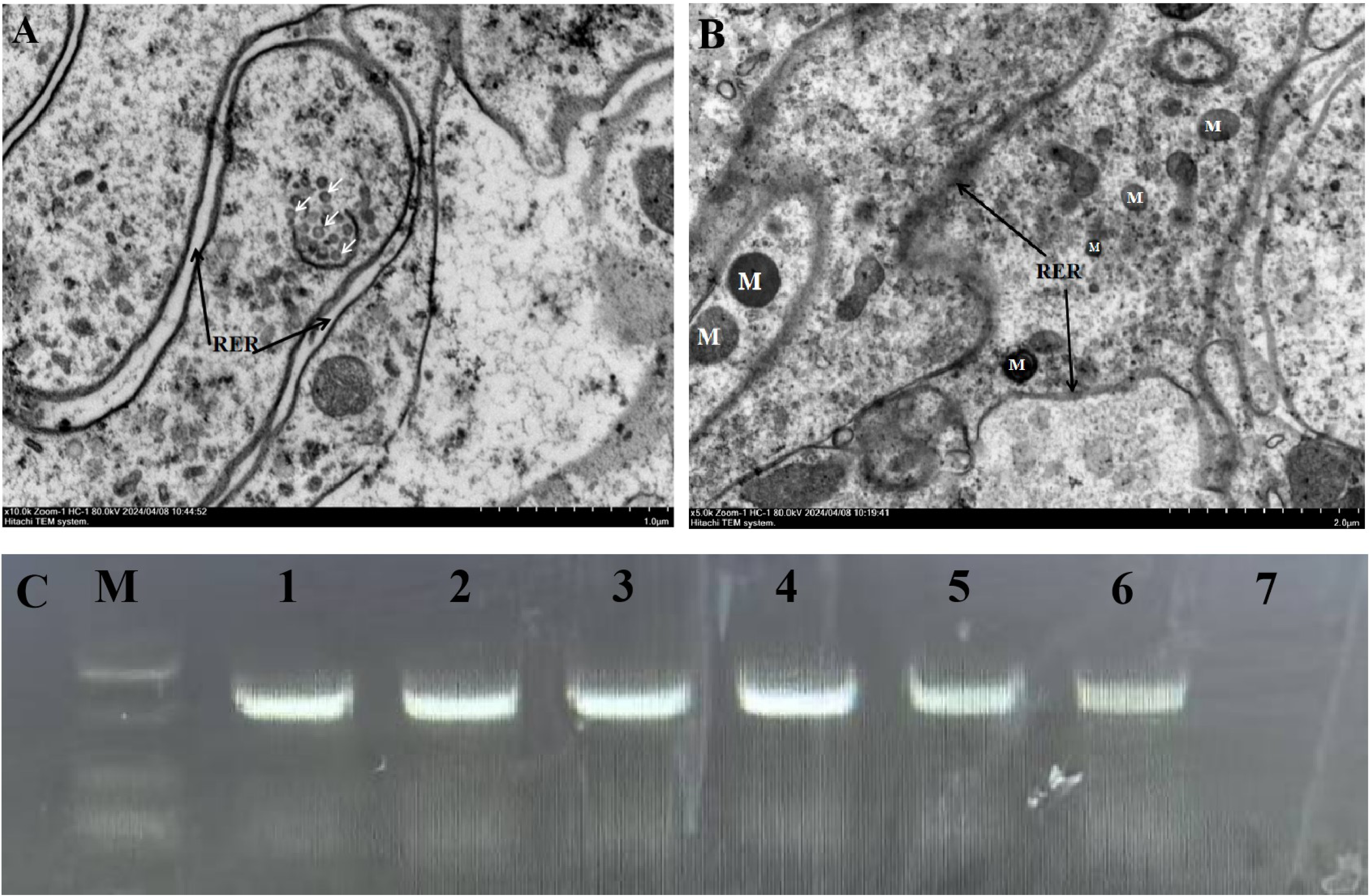
Detection of ASFV in different infected tissues of *O. lahorensis* adult ticks. **A:** Mature viruses emerged from the basement of the salivary glands of *O. lahorensis*, with proliferation of the rough endoplasmic reticulum (RER) by transmission electron microscopy. **B:** After infection, the mitochondria (M) in the tick salivary gland cells were closely associated with the factory and acquired a plasma membrane coating. **C:** PCR detection of ASFV in different infected tissues of *O. lahorensis* adult ticks. The “M”represents the 2000 molecular marker, The “1-6” numbers represent the salivary glands, midgut, Malpighian tubules, coxae, epidermis, and silk threads of infected ticks, respectively; The “7” represents the detection of uninfected ASFV ticks.

#### Detection of ASFV in ticks and pigs

To further confirm the ASFV load at different development stages of *O. lahorensis*, samples were collected for quantitative polymerase chain reaction (qPCR) analysis (Figure 4). The results revealed that B646L was expressed at the high levels at different developmental stages of the ticks. This process was accompanied by the development and sexual maturation of ticks and the viral content gradually increased. In this study, ticks that had already been infected with ASFV were allowed to develop (engorged larvae developed into hungry nymphs; engorged nymphs developed into hungry adults; engorged adults laid eggs), and hatched hungry larvae were used.

**Figure 4.**
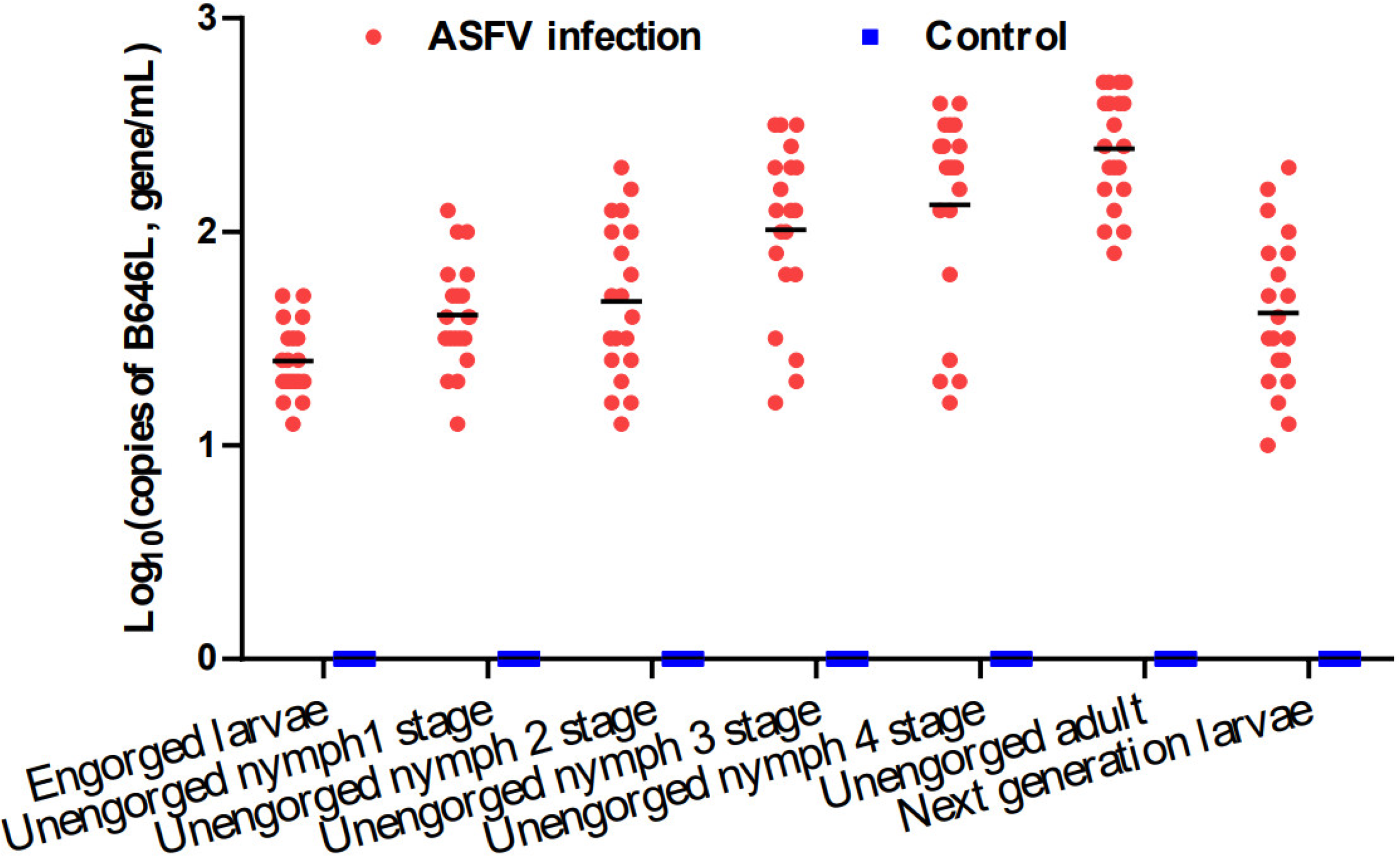
Detection of ASFV in *O. lahorensis* at different developmental stages. Each red dot represents one batch of sample (20 ticks).

These hungry ticks were randomly divided into three groups and released onto the skin of pigs that were not infected with ASFV. The viral content in the blood of the pigs and in the tissues of the dead pigs was measured every day. The results revealed that when engorged tick larvae developed into unengorged nymphs and bit clean experimental pigs, the amount of virus in the blood of the pigs gradually increased, with the highest content occurring at 9 ± 1 days. The content remained relatively stable at approximately 10 days (Figure 5B). In the spleen of the dead pigs at 11 days, the copy number of ASFV was relatively stable; the average ASFV content in the liver was 5.25 Log_10_ (copies of B646L, gene/mL), and that in the kidney was 4.79 Log_10_ (copies of B646L, gene/mL) (Figure 5A). Moreover, after the ASFV-infected nymphs developed into hungry adults, a similar trend as that described above was observed (Figure 5C). After the unengorged larvae bit the pigs, the ASFV content in the kidneys of pig #46 was greater than that in the spleen and liver, and the course of the disease in the pigs that died was longer (19±1 day) (Figure 5D).

**Figure 5.**
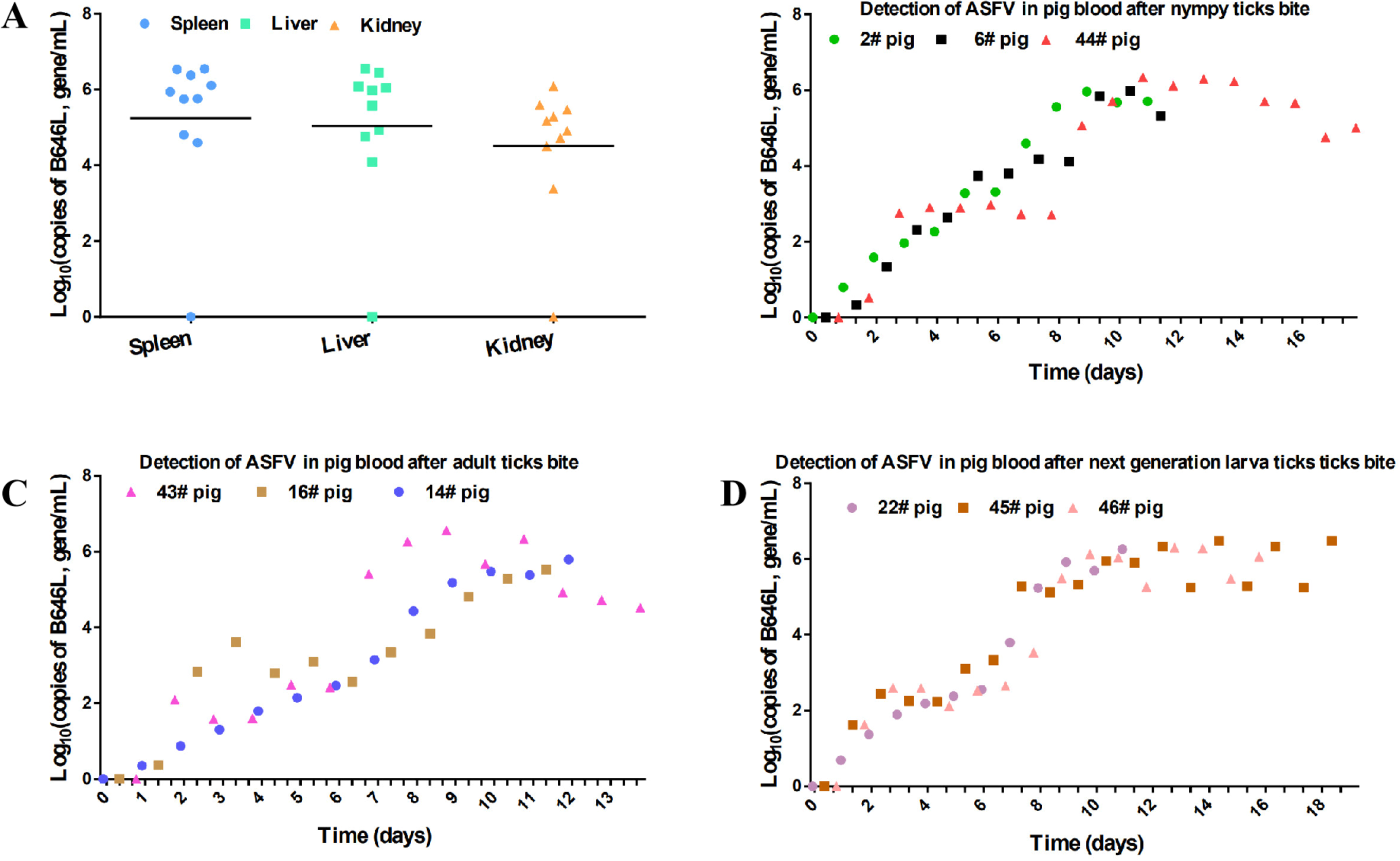
Detection of ASFV in the blood of pigs bitten by unengorged *O. lahorensis* ticks. **A**, Detection of ASFV in the spleen, liver, and kidneys of experimental pigs infected with ASFV via tick bites after pig death (uninfected tick-bitten pigs were used as controls). One colored dot in each sample represents a dead pig. **B**, Unengorged *O. lahorensis* nymphs infected with ASFV were randomly divided into three groups and allowed to bite some clean pigs. The #2 pig died on day 11. The #6 pig, whose tissue was collected on day 11, after death, with blood samples collected every day. The #44 pig, whose tissue was collected on day 18, after death, with blood samples were collected every day. **C**, After unengorged adult ticks infected with ASFV bit pigs #14, #16, and #43, daily testing of ASFV in pig blood was conducted. **D**, Detection of ASFV in the blood of pigs #22, #45, and #46 infected with ASFV after being bitten by larvae of the next generation.

## Discussion

The transmission dynamics of pathogens have garnered significant attention within the field of epidemiology, particularly with regard to vector-borne diseases. Recently, the confirmation that *O. lahorensis*, a soft tick species belonging to the family Argasidae, is capable of transmitting ASFV in China marked a substantial advancement in our understanding of the transmission pathways of this virulent pathogen. This discovery is particularly pertinent given the global implications of ASF, a disease that has wreaked havoc on pig populations worldwide, leading to economic devastation in sectors reliant on pig production ^31^. The aim of this section is to explore the implications, ecological considerations, and potential preventative strategies on the basis of the findings described herein, thereby providing a comprehensive overview of the current state of research surrounding this critical public health issue.

ASF is an acute viral disease that affects both domestic and wild pigs. The ramifications of ASF are particularly pronounced in China, which is home to the world’s largest pig population. Since the initial outbreak of ASF in China in 2018, the disease has led to unprecedented losses, eliminating more than one-third of the swine population of the country ^32^. The spread of ASF has far-reaching consequences not only for food security but also for the agricultural economy, employment, and the overall livelihood of farming communities ^33^. Consequently, the identification of novel vectors, such as *O. lahorensis*, underscores the need for a multidimensional approach for monitoring, controlling, and preventing this disease.

*O. lahorensis* is a soft tick species that primarily inhabits regions conducive to the life cycles of its host species, most notably the wild boar (*Sus scrofa*) ^34^. The epidemiological role of this tick species in facilitating the transmission of ASFV cannot be overstated. Research indicates that soft ticks tend to exhibit a prolonged feeding duration, which increases the likelihood of pathogen transmission during the process ^35^. Furthermore, their ability to thrive in diverse environmental conditions allows for sustained transmission dynamics, presenting a significant risk to pig populations ^36^. The confirmation of *O. lahorensis* as a competent vector for ASFV adds a layer of complexity to the already challenging landscape of disease management. Traditionally, control measures have focused on biosecurity protocols, vaccination efforts for susceptible domestic pig populations, and surveillance of known viral reservoirs ^37–39^. However, the vector-centric perspective that includes ticks as potential transmitters necessitates a comprehensive integration of acarology—the study of ticks—within these control strategies. This vector-borne transmission underscores the importance of ongoing surveillance not only of the virus but also of its vectors, expanding the remit of veterinary public health initiatives that aim to protect livestock. To fully understand the role of ticks as biological vectors of ASFV, we collected various types of ticks as close to the pig farm as possible, including hard and soft ticks of different genera. The test results revealed that none of the wild ticks tested were infected with ASFV. This finding indicates that ASFV has not entered wild ticks to form a natural focus. This finding is favorable for the prevention and control of ASFV. If the virus enters ticks, it will be difficult to eliminate, posing a major challenge for the prevention and control of ASFV.

Furthermore, there are significant public health implications associated with the recognition of ticks as vectors for ASFV. While ASF primarily impacts swine, its potential zoonotic nature, albeit not yet confirmed to directly affect humans, necessitates vigilance ^40^. Tick-borne pathogens have demonstrated the ability to adapt and evolve, posing risks of spillover into nontarget hosts, including humans. Therefore, comprehensive risk assessment protocols are needed to elucidate the broader epidemiological landscape surrounding ASF ^41^. In light of the alarming confirmation of *O. lahorensis* as a vector for ASFV, there is an urgent need for targeted interventions aimed at reducing tick populations and preventing their interactions with livestock. Integrated pest management approaches that encompass habitat modification, chemical control, and biological control strategies are imperative. Additionally, the development and implementation of effective acaricides are crucial for managing tick populations in areas at high risk for ASF outbreaks ^42^.

Vertical transmission refers to the passage of an infectious agent from one generation to the next, typically from parent to offspring. In the context of *O. lahorensis*, the concern lies in the ability of these ticks to harbor ASFV within their reproductive tissues and subsequently transmit it to their progeny. Studies have suggested that *O. lahorensis* can retain the virus through the transovarial transmission process, whereby the pathogen can infect the ovaries of female ticks, thus contaminating the eggs laid. The implications of vertical transmission are profound, as it enables ASFV to persist in the environment without necessitating continual reinfection from an external source. If female ticks can infect their offspring with ASFV, this complicates control efforts, as it potentially allows the establishment of reservoirs of the virus independent of infected pig populations ^43^. Understanding vertical transmission dynamics is critical for evaluating the role of *O. lahorensis* in the epidemiology of ASF and underscores the importance of tick control in managing this disease.

In contrast, transstadial transmission describes the capacity of a pathogen to endure within a tick as it advances through its life stages—from larva to nymph to adult. *O. lahorensis* experiences multiple developmental phases, during which it may consume blood from a variety of hosts. The importance of transstadial transmission in the context of ASFV is rooted in the virus’s potential to persist within these ticks throughout their life cycle, thus facilitating prolonged periods of infection. This mode of transmission presents significant risks ^44^, especially in settings where pigs are raised in close proximity to other animals or where ticks are permitted to move between species. The ability of *O. lahorensis* to carry ASFV throughout its life cycle implies that even if a pig is not directly infected, it could still be vulnerable to infection via interactions with ticks that harbor the virus.

TEM plays an important role as an extremely sophisticated observation tool ^45^. When studying ASFV in tick salivary glands, TEM allows detailed observation at the molecular level, enabling researchers to clearly identify the structure and layout of the virus within cells ^46,47^. These features are crucial for understanding how ASFV interacts with host tick cells and its lifecycle in such hosts. Through observation of tick salivary glands, researchers have shown that ASFV can form specific lesions within tick cells, typically manifesting as the aggregation of viral particles in the cytoplasm. Moreover, this study revealed the replication process of ASFV in tick salivary glands, where the virus invades host cells and utilizes the host’s biological mechanisms to reproduce extensively, thereby achieving transmission.

Studies have shown that ASFV is closely linked to the endoplasmic reticulum in tick salivary glands, suggesting that it utilizes the endoplasmic reticulum of cells as a site for replication. This discovery not only provides clues for understanding the lifecycle of ASFV but also lays the foundation for exploring its transmission pathways. However, despite providing important information for observing the mechanism of ASFV infection, TEM has limitations. A single observation result cannot represent all situations. By improving the understanding of the survival and transmission mechanism of ASFV in ticks, new ideas and methods can be provided for controlling the spread of the virus and reducing its impact on the pig industry. With continued technological advancement, we hope to see more innovative achievements in research in ticks and the pathogens they transmit.

## Conclusion

This study confirms that the *O. lahorensis* is another biological vector capable of transmitting ASFV. The identification of the *O. lahorensis* as a new vector means that pig farm management must become more comprehensive and meticulous. Traditional prevention and control measures have primarily focused on the isolation and culling of pigs, while neglecting the role of external parasites such as ticks. By enhancing monitoring and control of ticks, the risk of ASFV transmission can be effectively reduced, thereby protecting pig health and the sustainable development of the farming industry. Furthermore, this finding reminds us to strengthen our overall understanding of animal health. The spread of diseases is often the result of multiple factors. The inclusion of *O. lahorensis* highlights the need for farmers to consider various transmission vectors when implementing biosecurity measures, allowing for the development of more scientific and rigorous prevention strategies. This not only aids in the prevention and control of ASF but also helps improve the overall level of animal health management. Finally, recognizing the transmission capability of *O. lahorensis* calls for governments worldwide to enhance cross-border cooperation and information sharing, establishing a more comprehensive monitoring and response mechanism. Such global cooperation is essential for swiftly addressing outbreaks and minimizing impacts on international trade and food safety.

## Author Contributions

GYL and JL designed this study and critically revised the manuscript. HY, JXL and JL participated in its design, coordination. SYZ, QLN, JFY and QYR participated in sample collection. JL, Kashif, HJJ and GQG performed the experiments and data analysis and drafted the manuscript. All authors read and approved the final manuscript.

## Funding

This study was financially supported by the National Key Research and Development Program of China (2021YFD1800101); the Natural Science Foundation of Gansu Province (25JRRA437); the National Parasitic Resources Center, the Ministry of Science and Technology fund (NPRC-2019-194-30), the National Natural Science Foundation of China (U24A20450); the Key Program of the Nature Science Foundation of Gansu Province (25JRRA436); NBCITS (CARS-37) and Innovation Program of Chinese Academy of Agricultural Sciences (CAAS-ASTIP-2021-LVRI)

## Data Availability Statement

The data supporting the conclusions in this study are included in the article.

## Acknowledgments

All listed authors made a substantial, direct and intellectual contribution to this work and approved its publication.

## Conflicts of Interest

The authors declare that they have no conflflict of interest.

